# A vesicle-based platform for high-efficiency, high-viability CRISPR/Cas9 knockout in primary human myeloid cells

**DOI:** 10.64898/2025.12.28.696771

**Authors:** Silvia Fiori, Maira Russo, Alice Randon, Alessio Nicola Ferraro, Tiziana Julia Nadjeschda Schmidt, Elena Pinca, Rana K. Elnady, Ivan Castello Serrano, Paola Capasso, Abeer Mahfoud, Giorgia Moschetti, Edoardo Scarpa, Loris Rizzello, Niklas A. Schmacke, Veit Hornung, Takanobu Tagawa, Guido Papa, Manuel Albanese

## Abstract

Myeloid cells, including monocytes, macrophages, and dendritic cells, are central to host defence, inflammation, and antigen presentation. However, functional genetics studies in corresponding primary cells remain limited due to inefficient delivery methods. Here we report a vesicle-based CRISPR/Cas9 platform for effective genome editing in primary human myeloid cells using engineered virus-like particles (VLPs) and extracellular vesicles (EVs) to deliver Cas9-gRNA ribonucleoproteins (RNPs). Targeting primary CD14^+^ monocytes enables rapid generation of robust polyclonal knockouts while preserving cell viability and subsequent differentiation into macrophages and dendritic cells. Because our workflow relies on standard cell-culture handling rather than electroporation instrumentation, it is inexpensive, highly scalable and readily transferable across laboratories. Edited cells remain compatible with diverse downstream assays and functional readouts, enabling scalable loss-of-function studies and mechanistic dissection of primary human myeloid cell biology in health and disease. Vesicle-mediated RNP delivery thus provides a broadly applicable route to genetic perturbation in primary human myeloid lineages.

**Summary:** Fiori and colleagues describe a vesicle-based platform for efficient CRISPR/Cas9 knockout in primary human myeloid cells. This approach preserves cell viability, enables near-complete gene disruption, and supports functional studies of innate immunity, antigen presentation, and HIV-1 infection.

## Introduction

Macrophages are central sentinel cells of the innate immune system, connecting pathogen sensing to effector programs such as phagocytosis, inflammatory cytokine production, and antimicrobial responses (Park et al., 2022; Sweet et al., 2025; Wynn et al., 2013). Beyond their role in innate immunity, macrophages shape adaptive immune responses by processing and presenting antigens to T cells. They also support tissue homeostasis by clearing apoptotic cells and debris, remodeling tissue microenvironments, and coordinating repair and resolution of inflammation (Lazarov et al., 2023; Murray and Wynn, 2011; Sheu and Hoffmann, 2022; Watanabe et al., 2019). Despite this, many pathogens, including viruses and bacteria, infect, reprogram or exploit macrophages, turning them into reservoirs, vehicles for dissemination, or hubs of immune dysregulation (Cambier et al., 2014; Ferreira et al., 2024).

A major limitation in the study of human macrophages is the absence of experimental systems that combine physiological relevance with genetic tractability. Most studies rely on myeloid cell lines that are differentiated or polarized into macrophage-like cells (Schmacke et al., 2022), which only partially recapitulate primary macrophage diversity and plasticity. As a consequence, primary human macrophages represent a uniquely relevant system to dissect host-pathogen interactions and to define causal regulators of innate and adaptive immune circuits (Park et al., 2022).

Primary human monocytes can be isolated from peripheral blood as CD14^+^ cells. Although monocytes cannot be maintained *ex vivo*, they can be differentiated into monocyte-derived cells, including monocyte-derived macrophages (MDMs), inflammatory or anti-inflammatory macrophages, and monocyte-derived dendritic cells (moDCs), depending on the cytokine cocktail provided during differentiation (Martinez et al., 2006). Furthermore, primary human macrophages are terminally differentiated, non-proliferating and difficult to genetically manipulate, with limited lifespan in culture unless appropriately stimulated. Moreover, myeloid cells are highly sensitive to exogenous nucleic acids and other pathogen-associated molecular patterns (PAMPs), which can trigger innate sensing pathways and contribute to reduced delivery and editing efficiencies (Thompson et al., 2011). Indeed, genome editing of primary monocytes can be particularly challenging due to their response to nucleic acids and resistance to transduction (Gaidt et al., 2017; Greulich et al., 2019). Different solutions have been proposed to overcome these limitations, for example incorporating the HIV-2 accessory viral protein X (Vpx) into the delivered lentiviral vectors to counteract SAM and HD domain containing deoxynucleoside triphosphate triphosphohydrolase 1 (SAMHD1)-mediated restriction and enhance transduction efficiency (Hrecka et al., 2011), or directly deliver *in vitro*-preassembled CRISPR/Cas9-guide RNA (gRNA) ribonucleoproteins (RNPs) by nucleofection (Albanese et al., 2024; Freund et al., 2020; Hiatt et al., 2021). However, nucleofection, the most commonly used delivery method, induces considerable cellular stress and can reduce cell viability, while also requiring dedicated instrumentation, increasing experimental costs, and limiting scalability (Jung et al., 2025). In previous studies, nucleofection enabled KO efficiencies above 80%, but efficiency varied substantially between target genes (Freund et al., 2020; Hiatt et al., 2021). Since monocytes are quiescent and cannot be clonally expanded after editing, very high editing efficiencies are often required to reliably assess gene function in bulk populations. Here we present a vesicle-based CRISPR/Cas9 RNP delivery strategy that enables high editing rates in primary human myeloid cells while preserving viability and differentiation potential. We package and deliver Cas9 RNPs using engineered virus-like particles (VLPs) and extracellular vesicles (EVs), which protect their cargo and can be tailored for efficient protein transfer. By editing primary human monocytes prior to differentiation into macrophages, we generate polyclonal knockout populations with high reproducibility and subsequently obtain genetically perturbed macrophages and dendritic cells that are well suited for functional interrogation. Importantly, this approach is highly scalable and broadly accessible because it does not rely on specialized electroporation devices, thereby lowering the technical barrier for genetic manipulation of primary human myeloid cells. Together, these findings establish vesicle-mediated Cas9 RNP delivery as a simple, scalable, and efficient platform for loss-of-function studies across multiple targets in primary human myeloid cells, with broad applicability to downstream studies in innate and adaptive immunity, as well as viral infection models.

## Results

### MLV-based VLPs and EVs efficiently deliver CRISPR/Cas9 in human primary monocytes with comparable efficiency and viability to nucleofection

To establish a vesicle-based alternative to nucleofection, we generated engineered VLPs and EVs designed to deliver CRISPR/Cas9 RNPs into primary human CD14^+^ monocytes (Fig. 1 A). Both vesicle types are generated from HEK293T producer cells by transient transfection to express the components required for Cas9 RNP delivery, including a membrane-targeted Cas9 cargo (Cas9 fused to the viral structural protein Gag) and the vesicular stomatitis virus glycoprotein (VSV-G) as fusogenic envelope protein, which promotes target-cell entry and endosomal escape (Fig. 1 A). Freshly isolated human CD14^+^ monocytes are treated with the vesicles for target gene knockout, followed by knockout validation and functional phenotypic assays (Fig. 1 A).

**Figure 1.**
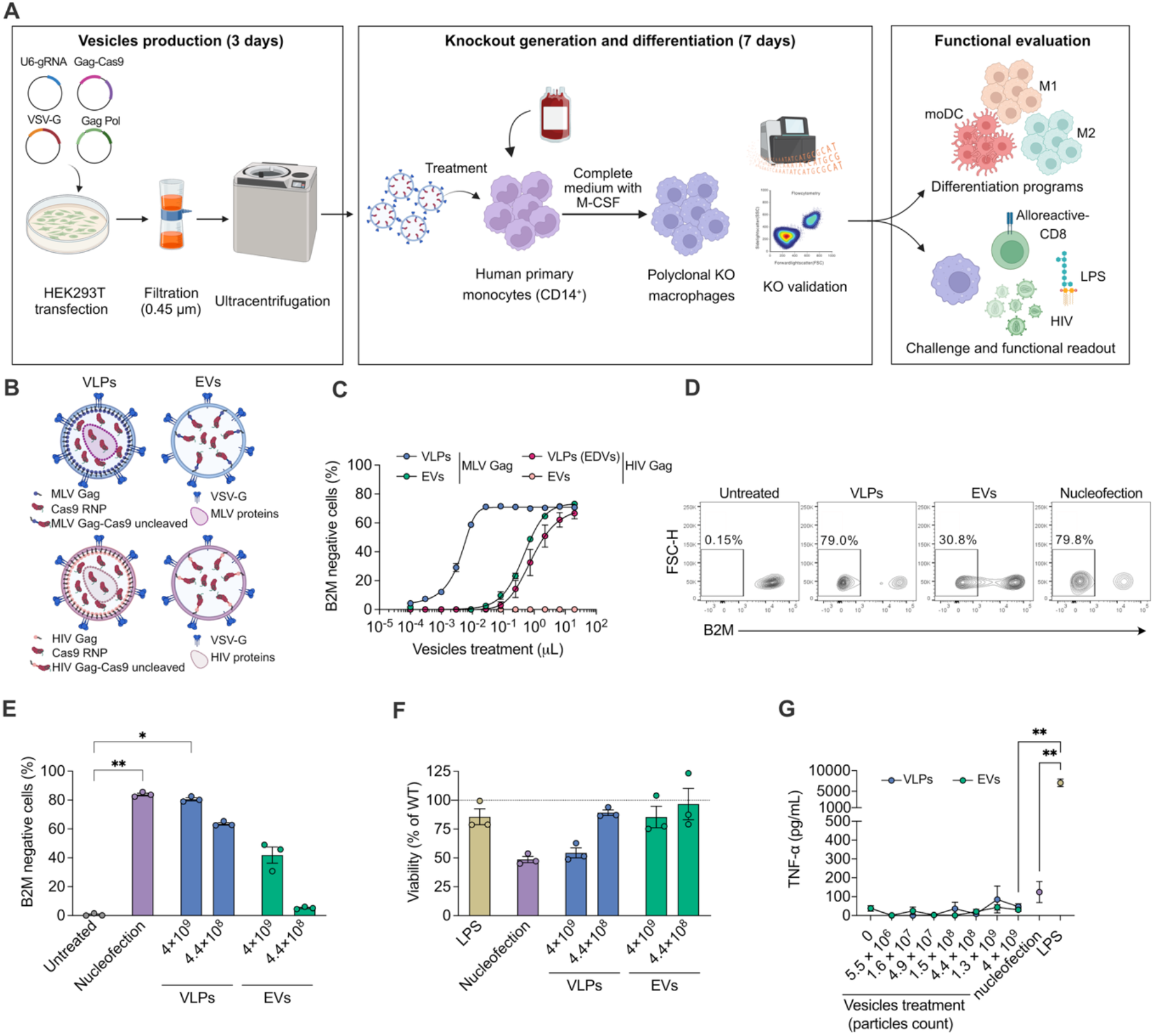
MLV-based vesicles efficiently deliver CRISPR/Cas9 in human primary CD14^+^ monocytes. **(A)** Schematic representation of the workflow for VLPs and EVs production, human primary monocytes (CD14^+^) treatment, monocytes differentiation in MDM, M1, M2, or moDC, knockout validation, and downstream functional characterizations. **(B)** VSV-G pseudotyped VLPs and EVs produced with MLV or HIV Gag polyproteins. The Cas9 is fused to the Gag polyprotein via a linker that is cleaved by the virus protease present in VLPs upon particle maturation. (**C**) B2M knockout efficiency in SUP-T1 cell line treated with increasing concentrations of EVs or VLPs carrying Cas9-RNP, where Cas9 protein is fused to MLV Gag or HIV Gag. B2M expression was measured by flow cytometry 7 days post-treatment (mean ± SEM; n=3). Representative flow cytometry plots (**D**) and bar plot (**E**) showing B2M knockout efficiency in MDM treated with increasing concentrations of EVs or VLPs carrying Cas9 fused to MLV Gag, compared with nucleofection with Cas9-gRNA RNP. B2M expression was measured by flow cytometry 7 days post-treatment (mean ± SEM; n=3 donors). Statistics indicate significance by Friedman test (* p < 0.05, ** p < 0.01). (**F**) Cell viability of differentiated MDM following EV/VLP treatment, nucleofection (as in **E**) or LPS as control, assessed by CellTiter-Glo (mean ± SEM; n=3 donors). (**G**) TNF-α production by MDMs after EVs and VLPs treatment, nucleofection, or LPS stimulation (positive control), quantified by ELISA on supernatants collected after overnight incubation (mean ± SEM; n=3 donors). Statistics indicate significance by two-way ANOVA. P values were corrected for multiple comparison (Tukey) (* p < 0.05, ** p < 0.01).

The principal distinction between the two formulations is that EVs are produced without the additional viral packaging/structural components used to assemble VLPs (Fig. 1 B). This feature is expected to reduce the overall viral protein content of the particles but may also necessitate additional optimization to maximize Cas9 incorporation, as Gag-driven assembly and cargo loading are intrinsically more efficient in VLPs. Two VLPs-based platforms are currently the most widely used for Cas9-based gene editor delivery: Murine Leukemia Virus (MLV)-based VLPs (Banskota et al., 2022) and human immunodeficiency virus (HIV)-based Cas9 Enveloped Delivery Vehicles (Cas9EDVs)(Hamilton et al., 2024). We compared the ability of these two VLP systems by loading them with MLV Gag-Cas9 or HIV Gag-Cas9, respectively, together with a single gRNA targeting the β2-microglobulin (*B2M*) gene (Fig. 1 C) in the SUP-T1 cell line, given its high sensitivity to VSV-G-mediated entry. In parallel, we also produced EVs containing either MLV Gag-Cas9 or HIV Gag-Cas9. Interestingly, MLV-based VLPs and EVs achieved higher B2M KO levels compared to HIV-based vesicles (∼30-fold higher for MLV-VLPs versus HIV-VLPs) (Fig. 1 C). For this reason, we exploited MLV Gag-based vesicles as the delivery tool for Cas9 RNPs in myeloid cells. First, we assessed VLPs loaded with a *B2M*-targeting single gRNA in freshly isolated primary monocytes achieving KO efficiencies comparable to RNP nucleofection (Fig. 1 D,E). EVs delivery of B2M-targeting single gRNA reached high KO efficiencies, yet it was less efficient than VLPs or nucleofection (Fig. 1 E). Analyzing the targeted locus directly by deep sequencing (Illumina Miseq) confirmed high genome editing rates (Fig. S1 A,B).

Surprisingly, we observed a loss of viability at high VLPs concentrations comparable to that observed after nucleofection (Fig. 1 F), even though we did not observe increased pro-inflammatory responses (TNF-α) (Fig. 1 G). We reasoned that this loss of viability could be due to residual impurities in the vesicle preparations. To test this hypothesis, we used alternative purification methods, recommended by the International Society for Extracellular Vesicles (ISEV) guidelines (Welsh et al., 2024). We isolated VLPs from the cell supernatant by ultracentrifugation (our standard procedure), followed by an additional purification step using either density gradient (DG) separation or size-exclusion chromatography (SEC)(Albanese et al., 2021). The different isolation methods achieved comparable KO efficiencies (Fig. S1 C), but none improved cell viability after treatment (Fig. S1 D). Notably, exposure of monocytes to ultracentrifugation-purified VLPs lacking the vesicular stomatitis virus glycoprotein (VSV-G) did not reduce cell viability and, as expected, failed to induce KO (Fig. S1 C,D). This is consistent with prior observations that non-pseudotyped (“naked”) vesicles do not efficiently deliver their cargo to target cells (Albanese et al., 2021; An et al., 2024; Banskota et al., 2022; de Jong et al., 2020; Somiya and Kuroda, 2021a). Together, these data suggest that reduced fitness is unlikely to be driven by vesicle impurities, but rather by efficient cargo delivery and/or VSV-G itself.

### Low doses VSV-G vesicles are highly functional and preserve macrophage viability

VSV-G is a widely utilized envelope protein for pseudotyping both viral and non-viral vectors. It mediates particle binding to low-density lipoprotein receptors (LDLR) on target cells, facilitating subsequent endosomal escape. Beyond entry kinetics, incorporation of VSV-G into extracellular vesicles has been reported to enhance vesicle release from producer cells (Bui et al., 2023; Zhang et al., 2020). However, high or sustained VSV-G expression can be cytotoxic to producer cells. This trade-off prompted us to examine how VSV-G incorporation into vesicles affects both genome editing efficiency and recipient cell viability. To identify the optimal VSV-G dose for our platform, we generated VLPs using increasing amounts of VSV-G plasmid during producer-cell transfection, starting from our standard 10% formulation, defined as the percentage of total transfected DNA, and tested their activity in SUP-T1 cells. We found that a 20-fold reduction of VSV-G from 10% (VSV-G high) to 0.5% (VSV-G low) did not affect genome editing efficiency (Fig. 2 A). However, VSV-G low VLPs led to a marked increase in SUP-T1 cell viability following VLPs treatment (Fig. 2 B). Consistent with these findings, treatment of primary macrophages with VSV-G low VLPs similarly improved cell fitness and morphology while preserving KO efficiencies comparable to VSV-G high conditions (Fig. 2 C,D).

**Figure 2.**
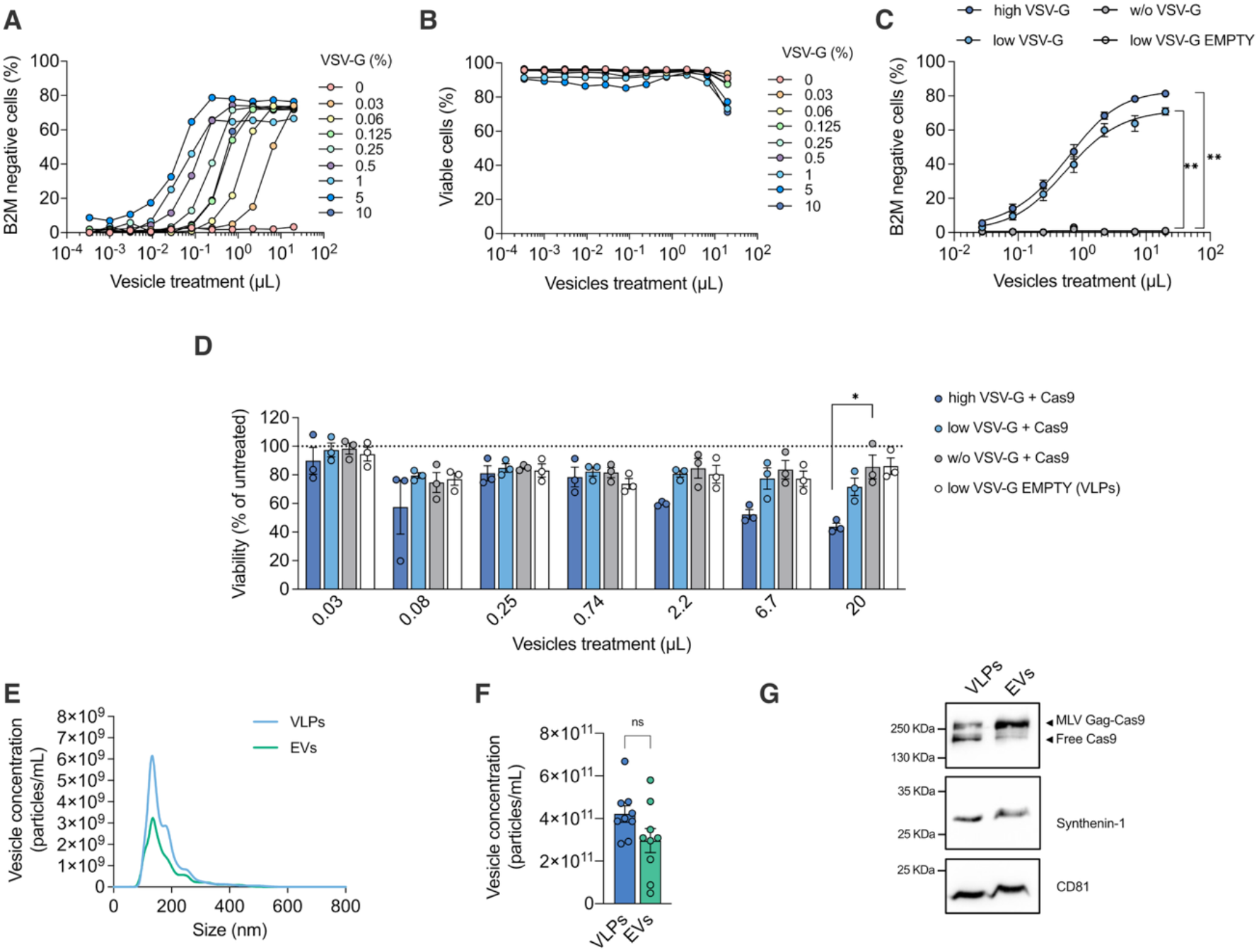
Reduced VSV-G content improves macrophage viability while preserving vesicle structure and function. (**A**) B2M knockout efficiency in SUP-T1 cell line treated with equal volumes of EVs produced with decreasing VSV-G levels. The percentage of VSV-G indicates the fraction of the VSV-G plasmid relative to the total DNA used for transfection. B2M expression was analysed by flow cytometry 5 days post-treatment. One representative experiment is shown (n=3). (**B**) Viability analysis measured by flow cytometry for the experiment shown in **A**. One representative experiment is shown (n=3). (**C**) B2M knockout efficiency in primary MDMs assessed by flow cytometry 7 days after treatment with VLPs high (10%) or low VSV-G (0.5%) (mean ± SEM; n=3 donors). Statistics indicate significance by two-way ANOVA. P values were corrected for multiple comparison (Tukey) (** p < 0.01). (**D**) Cell viability relative to the experiment shown in **C** (mean ± SEM; n=3 donors). Statistics indicate significance by two-way ANOVA. P values were corrected for multiple comparison (Tukey) (* p < 0.05). (**E**) Size distribution profiles of EVs and VLPs measured by NTA after ultracentrifuge-based purification. Mean of three independent VLPs and EVs particles (mean ± SEM; n=3 independent productions). (**F**) Particle concentration after ultracentrifugation-based purification (mean ± SEM; n=9). Statistic indicates not significance by unpaired t-test. (**G**) Cas9 detection in EVs, and VLPs by western blot. Free Cas9 and MLV Gag-Cas9 fusion protein are indicated with black arrows. One representative blot is shown (n=2).

Interestingly, decreasing VSV-G levels reduced particle concentration exclusively in EVs formulations (Fig. S2 A). In contrast, VLPs remained unaffected, likely because MLV packaging proteins dominantly drive particle assembly and release, overriding variations in VSV-G content (Fig. S2 A,B).

VSV-G-low VLPs and EVs showed a relatively uniform size distribution, with an average particle size of approximately 150 nm, and EVs preparations displayed slightly lower particle concentrations than VLPs preparations, despite some degree of variability (Fig. 2 E,F and Fig. S2 C). Western blot analysis revealed efficient Cas9 loading within both vesicle types. However, whereas the cleaved, free form of Cas9 was predominant in VLPs due to MLV protease-mediated processing of the Gag-Cas9 fusion protein (Banskota et al., 2022), EVs lacked this viral protease and primarily retained the intact, uncleaved Gag-Cas9 protein (Fig. 2 G). Collectively, these findings allowed us to identify a robust vesicle-based platform based on low VSV-G content for high-efficiency genome editing in primary monocytes while preserving cell viability overcoming key limitations associated with nucleofection.

### Low VSV-G vesicles enable complete knockout of the target protein and preserve monocyte differentiation programs

Achieving complete knockout of a target gene is particularly important in primary monocyte-derived systems, as it allows edited cells to be maintained as polyclonal populations and avoids the need for single-cell cloning or FACS-based sorting, both of which are technically challenging and can impose substantial stress on differentiated macrophages. Since increasing VLP or EV doses did not raise knockout efficiencies above 80% with a single gRNA, we tested a second gRNA targeting the same locus (gRNA2) and the combined delivery of both guides (gRNA 1+2) (Fig. 3 A and Fig. S2 D). Co-delivery of the two gRNAs substantially enhanced editing efficiency, achieving a complete KO in monocytes differentiated into MDMs (Fig. 3 B) or into monocyte-derived dendritic cells (moDCs) (Fig. 3 C), while preserving also moDCs viability (Fig. 3 D). EVs treatment also reached a complete B2M KO, although ∼8-fold more particles were required to achieve similar levels to VLPs, likely due to higher presence of free-Cas9 form in VLPs.

**Figure 3.**
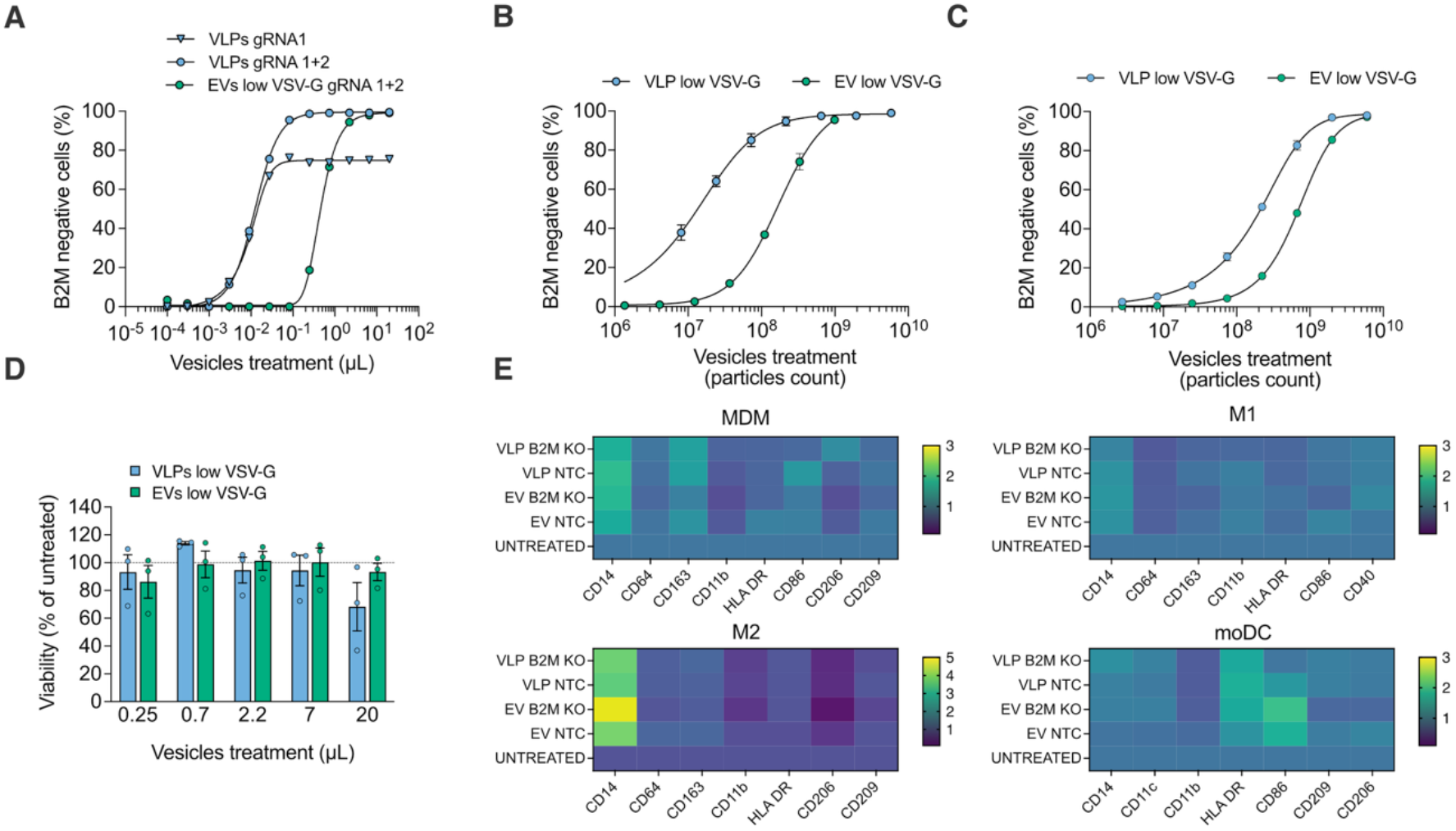
Vesicle-mediated B2M knockout in monocyte-derived macrophages and dendritic cells preserves viability and differentiation program. (**A**) B2M knockout in SUP-T1 cells assessed by flow cytometry 5 days after treatment, comparing a single gRNA versus a two-gRNAs combination loaded in VLPs and EVs. (**B**) B2M knockout efficiency in MDMs treated with increasing dose of EVs or VLPs, VSV-G low incorporating 2 gRNAs together. B2M expression was measured by flow cytometry 7 days post-treatment, results were normalized to particle number (mean ± SEM; n=3 donors). (**C**) B2M knockout efficiency in moDCs treated with increasing concentrations of EVs or VLPs, VSV-G low incorporating 2 gRNAs. B2M expression was measured by flow cytometry 7 days post-treatment, results were normalized to particle number (mean ± SEM; n=3 donors). (**D**) Cell viability of moDCs following EVs/VLPs treatment (as in C), assessed by CellTiter-Glo (mean ± SEM; n=3 donors). Statistical analysis indicate not significance by Wilcoxon test. (**E**) Differentiation markers that identifies MDM, M1, M2, and moDC populations. Monocytes were treated with VLPs and EVs targeting B2M or incorporating non-targeting control (NTC) gRNA followed by differentiation with different cytokine combinations (see material and methods). The heatmap shows fold change values of Median fluorescence intensity (MFI) with respect to untreated control (set as 1), measured by flow cytometry (mean ± SEM; n=3 donors).

Primary monocytes can differentiate *in vitro* into unpolarized (MDM) macrophages upon M-CSF stimulation. Subsequent exposure to specific cytokine cocktails polarizes these cells toward pro-inflammatory or immunosuppressive phenotypes, which are conventionally referred to M1 and M2 (Mantovani et al., 2004; Murray, 2017; Wynn and Vannella, 2016). Beyond macrophage lineage commitment, monocyte developmental plasticity allows for differentiation into moDCs, which are often exploited for their antigen presenting function in several *in vitro* or *ex vivo* assays(Sallusto and Lanzavecchia, 1994). To demonstrate that monocytes treated with VLPs and EVs retain their polarization potential, we differentiated monocytes into MDM, M1, M2, and moDC after treatment with VLPs and EVs incorporating two gRNAs targeting B2M gene or with incorporating a non-targeting control (NTC) gRNA and comparing these conditions with untreated cells (Fig. 3 E and Fig. S3). No significant differences were observed in the canonical markers defining each polarization state. However, M2 macrophages treated with VLPs or EVs retained higher CD14 expression compared with untreated controls, along with a modest, non-significant reduction in CD206 (Fig. 3 E). Additionally, treatment of moDCs with both particle types slightly upregulated CD86 and HLA-DR expression, suggesting that VLPs and EVs partially promote moDC maturation and activation. Together, these findings show that vesicle-mediated dual-gRNA delivery achieves complete polyclonal knockout without compromising the developmental plasticity of primary human monocytes, supporting its use for functional studies across macrophage and dendritic-cell differentiation states.

### Vesicle-based knockout enables functional gene interrogation in primary human macrophages

Macrophages and dendritic cells are key players in immune responses, acting as a bridge between innate and adaptive immunity. They are also targeted by diverse pathogens, including HIV, mycobacterium tuberculosis, and CMV (Baasch et al., 2021; Cambier et al., 2014; Ferreira et al., 2024). In this context, genetic perturbation represents a powerful tool to dissect the functional pathways involved in immune regulation and host-pathogen interactions. To determine whether macrophages edited using our vesicle-based platform remain suitable for downstream functional readouts, we knocked out, as proof-of-concept examples, three key genes involved in distinct biological contexts: B2M for adaptive immunity, TLR4 for innate immune signaling, and CCR5 for host-pathogen interaction.

To assess whether vesicle-edited macrophages could be used for functional antigen-presentation assays, we generated B2M-KO MDMs and, in parallel, MDMs treated with vesicles carrying a NTC gRNA.

Because B2M is required for HLA-I surface expression, HLA-I-restricted recognition by CD8 T cells provides a direct and biologically relevant functional readout of B2M knockout (Fig. 4 A).

**Figure 4.**
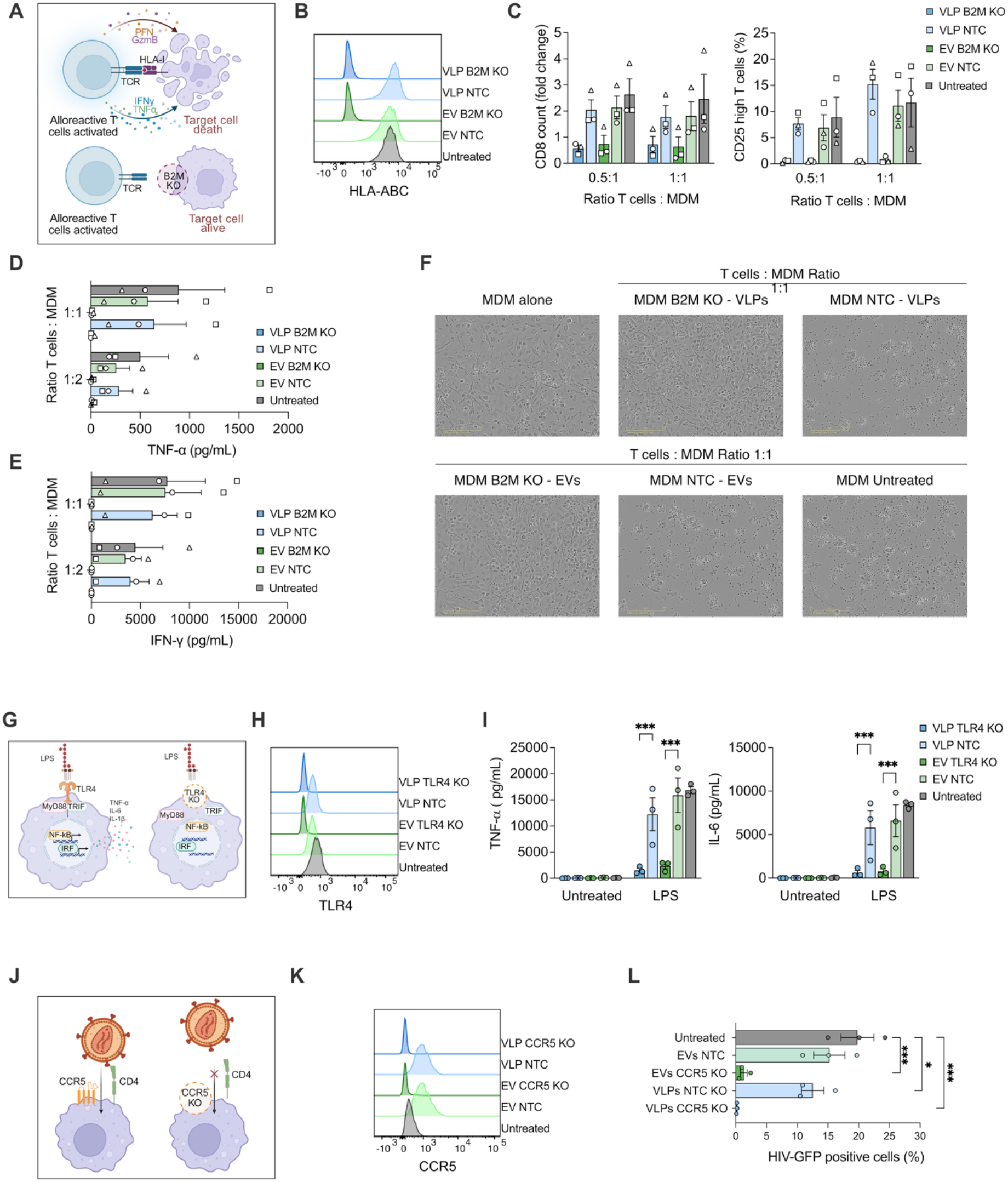
Vesicle-mediated B2M knockout in monocyte-derived macrophages and dendritic cells supports immunological and virological downstream functional assays. (**A**) Schematic of the co-culture setting used to assess functional outcomes of B2M knockout. (**B**) HLA-ABC expression in MDM treated with VLPs and EVs targeting B2M or with NTC gRNA control measured by flow cytometry. One representative experiment is shown (n=3). (**C**) Percentage of number CD8 (left) and CD25 positive cells (right) measured by flow cytometry after 7 days from co-culturing alloreactive CD8 T cells specific for HLA-A02 alleles and MDM from HLA-A02 positive donors treated with VLPs and EVs targeting B2M or with NTC gRNA or untreated. Ratios of T cells to MDM was 0.5:1 and 1:1 (mean ± SEM; n=3 donors). (**D)** TNF-α and (**E**) IFN-γ overnight secretion in co-culture supernatants (as in C), quantified by ELISA (n=3). (**F**) Qualitative assessment of target-cell killing by microscopy of alloreactive CD8 T cells in co-cultures with edited MDMs (B2M KO) or NTC (T:MDM ratio 1:1), compared with vesicle-untreated controls. Images shown are from one representative donor experiment (n=2). (**G**) Schematic representation of the functional readout setting of edited MDMs (TLR4 KO) or MDM NTC stimulated with LPS. (**H**) TLR4 KO efficiency in MDMs treated with increasing concentrations of EVs or VLPs VSV-G low incorporating 2 gRNAs together. TLR4 expression was measured by flow cytometry 7 days post-treatment. One representative experiment is shown (n=3). (**I**) TNF-α and IL-6 released by MDMs treated as explained in **G** upon 6 hour-stimulation with LPS (mean ± SEM; n=3 donors). (**J**) Schematic representation of HIV infection treatment in edited MDMs and NTC MDMs. (**K**) CCR5 KO efficiency in MDMs treated with increasing concentrations of EVs or VLPs VSV-G low incorporating 2 gRNAs together. CCR5 expression was measured by flow cytometry 7 days post-treatment. One representative experiment is shown (n=3). (**L**) HIV infection percentage in CCR5 KO, NTC, and untreated MDMs measured by FACS 3 days post infection (mean ± SEM; n=3 donors).

We first confirmed that B2M knockout reduced HLA-I surface expression on MDMs (Fig. 4 B). We then assessed whether B2M-deficient MDMs were impaired in their capacity to be recognized by alloreactive CD8 T cells. To generate HLA-I-specific alloreactive effectors, CD8 T cells from an HLA-A*02-negative donor were repeatedly stimulated with HLA-A*02-positive feeder cells, yielding polyclonal CD8 T-cell lines reactive against HLA-A*02-expressing targets.We co-cultured these alloreactive CD8 T cells with untreated, NTC-treated, or B2M-KO MDMs as target cells. In this setting, CD8 T cells proliferated and became activated when co-cultured with untreated or NTC-treated MDMs, but not with B2M-KO MDMs, as measured by CD8 T-cell counts (Fig. 4 C left) and CD25 expression (Fig. 4 C right). Measurement of IFN-γ and TNF-α secretion by ELISA further corroborated these results (Fig. 4 D,E).

Finally, to evaluate the durability of vesicle-edited MDMs and their suitability for longer-term functional assays, we extended the co-culture to 7 days using a high effector-to-target ratio of 1:1. Even under these stringent conditions, B2M-KO MDMs remained viable after one week, whereas untreated and NTC-treated MDMs were efficiently eliminated by alloreactive CD8 T cells (Fig. 4 F).

Having established that vesicle-edited MDMs are suitable for functional assays of antigen presentation and CD8 T-cell recognition, we next tested whether the same platform could be used to interrogate innate immune signalling. To this end, we targeted TLR4, the canonical receptor for lipopolysaccharide (LPS), which drives pro-inflammatory cytokine secretion through MyD88-and TRIF-dependent pathways (Akira et al., 2006; Thompson et al., 2011)(Fig. 4 G). Freshly isolated monocytes were treated with EVs or VLPs loaded with either dual *TLR4*-targeting gRNAs or NTC. Both delivery vehicles achieved complete TLR4 protein ablation (Fig. 4 H). Following 7 days of culture with M-CSF, the differentiated MDMs were challenged with 50 ng/mL LPS. While untreated and NTC-treated MDMs mounted a robust inflammatory response characterized by IL-6 and TNF-α secretion, this cytokine production was abolished in TLR4-KO MDMs (Fig. 4 I).

As a third proof-of-concept application, we targeted the CCR5 chemokine receptor, which is expressed on CD4 T cells and macrophages and serves as a major co-receptor for HIV-1 entry (Alkhatib et al., 1996; Deng et al., 1996)(Fig. 4 J). We first verified that CCR5 was efficiently knocked out using both VLPs and EVs (Fig. 4 K). Next, to maximize viral infection, we treated MDMs with VLPs carrying the viral protein Vpx to counteract SAMHD1, a major restriction factor for HIV-1 infection in myeloid cells (Hrecka et al., 2011), and then infected the cells with an R5-tropic HIV-1 GFP reporter virus (Albanese et al., 2022). As expected, CCR5-KO macrophages were protected from infection with the HIV-1-GFP reporter virus, whereas NTC-treated and untreated macrophages remained susceptible to viral infection (Fig. 4 L).

Together, these data establish vesicle-based RNP delivery as a straightforward, highly effective, and immunologically neutral strategy to edit primary monocytes while preserving their viability and differentiation capacity. Importantly, MDMs edited using this vesicle-based gene editing platform are directly compatible with downstream functional assays without the need for selection or sorting.

## Discussion

Primary human myeloid cells are central to host defence, inflammation, and antigen presentation, but they remain challenging models for functional genetics studies. Several groups have made important progress using CRISPR/Cas9 RNPs delivered by nucleofection to freshly isolated CD14 monocytes. This approach enables efficient gene knockout (Freund et al., 2020; Hiatt et al., 2021) and has opened the door to phenotype-driven studies that were previously inaccessible in primary human myeloid cells (Albanese et al., 2024).

However, nucleofection-based workflows induce substantial cellular stress, reduce cell viability, and are costly and difficult to scale, especially when near-complete knockout must be achieved across multiple targets, donors, or experimental conditions. These constraints become particularly restrictive for primary human monocytes, where cell numbers are often limiting and incomplete editing can compromise the interpretation of downstream functional assays performed in bulk populations. Here, we present a vesicle-mediated, DNA-free delivery of pre-assembled CRISPR/Cas9 RNPs that enables genome editing in primary human monocytes while preserving viability and differentiation capacity. We package pre-assembled Cas9-gRNA RNPs into engineered VLPs or EVs. These vesicle-based strategies enable transient protein delivery, thereby minimizing exposure to exogenous DNA and limiting prolonged nuclease expression. VLPs and EVs-mediated delivery yielded robust polyclonal knockout in primary monocytes and, importantly, edited monocytes could be differentiated into MDMs and moDCs without selection or sorting, enabling straightforward downstream functional assays.

A central technical insight from our work is that delivery efficiency and viability can be independently optimized by tuning the fusogenic component required for productive cargo transfer. Although additional purification (density gradient or size-exclusion chromatography) did not improve viability, VLPs produced without the fusogenic envelope protein VSV-G preserved viability but also failed to edit the genome of target cells, consistent with inefficient cytosolic cargo delivery by non-pseudotyped vesicles. This aligns with broader evidence that engineered, DNA-free vesicles can efficiently deliver RNP payloads, but that productive entry and cargo release often depend on envelope-mediated fusion and consequent endosomal escape (Albanese et al., 2021; Banskota et al., 2022; de Jong et al., 2020; Hamilton et al., 2021; Somiya and Kuroda, 2021b).

By reducing VSV-G during vesicles production, we identified a minimal level of fusogenic molecule that preserves editing potency while improving cell fitness across contexts, including primary macrophages. This practical titration strategy provides a simple lever to optimize delivery for sensitive primary myeloid cells, and it highlights fusogenicity as a dominant determinant of tolerability under otherwise comparable purification and dosing conditions.

Our data also emphasize the value of multiplexing to drive near-complete knockout in non-proliferating primary models. While a single guide achieved robust but incomplete KO of B2M, combining two gRNAs markedly increased KO efficiency to a complete phenotype across monocyte-derived lineages. This concept is consistent with reports that multiplexing gRNAs in optimized nucleofection workflows for primary myeloid cells can yield additive increases in knockout efficiency, thereby reducing the need for enrichment strategies that are impractical in terminally differentiated populations or difficult-to-culture cells (Albanese et al., 2022; Freund et al., 2020). Our results extend this principle to vesicle-based delivery. In this context, multiplexing offers a previously underappreciated advantage by enabling strong phenotypes at lower particle doses, improving cellular tolerability while maintaining functional knockout. B2M was selected as a benchmark because it is required for HLA class I surface stability and is ubiquitously expressed across nucleated cells. In a demanding co-culture system with alloreactive CD8 T cells, B2M-KO MDMs failed to induce T cell activation and expansion, as also confirmed by the absence of IFN-γ secretion. Importantly, under prolonged, high-pressure conditions (7-day co-culture at high effector-to-target ratio), B2M-KO macrophages were spared from cytotoxic CD8 T cells, whereas control macrophages were efficiently eliminated. Together, these results demonstrate that vesicle-based editing not only produces high knockout efficiencies, but also preserves expected cellular behavior in prolonged experimental setups, including differentiation state, functional competence, and long-term survival. The TLR4 and CCR5 experiments further extend this validation to innate immune signaling and host-pathogen interaction models, showing that vesicle-edited macrophages can support both acute stimulation assays and infection-based functional readouts. This distinguishes our approach from stress-inducing workflow and establishes vesicle-mediated delivery as a practical and tractable platform for phenotypic interrogation of antigen presentation, host-pathogen interactions, and immune effector function in primary human macrophages.

Our findings position vesicle-based delivery as a complementary approach to existing nucleofection-based platforms. Nucleofection workflows are rapid and can achieve high KO efficiencies across multiple myeloid contexts, including protocols that edit monocytes prior to differentiation while preserving key functional properties such as phagocytosis and marker expression. However, nucleofection frequently compromises cell viability, necessitates careful monitoring of activation states, and often depends on access to optimized protocols and electroporation instrumentation. Our vesicle-based approach mitigates these constraints by shifting delivery to a transient, DNA-free, non-electroporation modality implemented with standard cell-culture handling. Ready-to-use vesicles can be cryopreserved (-80 °C) (Gorgens et al., 2022) and added directly to plated monocytes to achieve complete knockout without specialized equipment, providing a practical advantage when viable yield, long-term culture, and differentiation fidelity are limiting.

Beyond this practical workflow, our approach builds on a growing body of work demonstrating that engineered, DNA-free VLPs can serve as efficient carriers for RNP payloads. Such systems have been successfully applied to deliver a range of CRISPR-based editors, including base and prime editors both *in vitro* and *in vivo* delivery contexts (An et al., 2024; Banskota et al., 2022; Geilenkeuser et al., 2025; Hamilton et al., 2024). Consistent with this framework, our MLV-based VLPs production strategy was adapted and optimized for myeloid cells from a previously described system (Banskota et al., 2022).

Here, we extend these principles to primary human myeloid editing and explicitly focus on the delivery-toxicity trade-off that emerges in monocytes and macrophages. Notably, recent studies have further expanded the VLP toolkit, enabling base editing, epigenetic perturbation, knock-in strategies, and even pooled screening approaches in primary myeloid cells, while maintaining low toxicity and preserved innate immune responsiveness (Jung et al., 2025). In this context, our data suggest that MLV-based VLPs achieve higher KO efficiency than HIV-based Cas9EDVs (Banskota et al., 2022; Hamilton et al., 2024; Jung et al., 2025). Additionally, we expand the available delivery platforms by including EVs, which achieved comparable KO efficiencies to VLPs, albeit at higher doses, while substantially reducing the contribution of viral proteins to the delivery system. Although EVs may not offer a major advantage for *in vitro* editing in our current setting, their reduced viral-protein content and potentially lower immunogenicity may make them attractive for future *in vivo* applications requiring repeated dosing (Tian et al., 2026). A practical limitation of vesicle-mediated delivery is the need for vesicle production and quality control before use. However, this workflow is based on standard transient transfection of HEK293T producer cells and simple concentration steps, making it readily accessible even to non-specialist laboratories. Future optimization of cargo loading, purification, and storage may further improve potency and reproducibility, particularly for EV-based formulations.

Taken together, our findings align with and extend this trajectory and motivate future applications of the platform beyond gene knockout. These include the use of targeted envelopes to improve cell-type selectivity, alternative fusogens to further optimize tolerability, and coupling to richer readouts such as single-cell profiling. Similar strategies have already been explored in other primary cell types, including T cells (Frank and Buchholz, 2019; Hamilton et al., 2024) and hematopoietic stem cells (Botchkarev et al., 2025), as well as *in vivo* delivery settings.

In summary, we present a vesicle-based CRISPR/Cas9 RNP delivery platform that enables high-efficiency, polyclonal genome editing in primary human monocytes while preserving viability and differentiation into macrophages and dendritic cells. By defining a practical fusogen-tuning strategy that improves tolerability without compromising delivery efficiency, and by demonstrating stable functional phenotypes in stringent co-culture assays, our work provides a broadly accessible framework for mechanistic studies of human myeloid biology. More broadly, this platform enables scalable loss-of-function studies across donors, accelerates causal mapping of innate and adaptive immune circuits, and supports disease-relevant applications in which myeloid cell fitness and functional integrity are essential.

## Materials and Methods

### Cell culture

HEK293T cells (DSMZ ACC 635) were maintained in complete DMEM supplemented with 10% fetal bovine serum (FBS), 1% penicillin–streptomycin, 1% non-essential amino acids, 1% L-glutamine and 1% sodium pyruvate (all Gibco). SUP-T1 cells (DSMZ ACC 140) were maintained in complete RPMI 1640 10% FBS, 1% penicillin–streptomycin, 1% non-essential amino acids and 1% sodium pyruvate (all Gibco). All cells were cultured at 37 °C with 5% CO₂ in a humidified incubator. FBS EV-depleted was used to produce VLPs and EV and obtained as previously described(Albanese et al., 2021). FCS was diluted 1:1 with medium and centrifuged at 100,000 g at 4°C in a swinging bucket rotor (SW32, Beckman Coulter) for 18 h. The supernatant was filter and sterilized using a 0.22-μm mesh size filter (Sartorius) and then filtered using a 300K Vivaspin 20 (PES, Sartorius) device at 2,000 g at 10°C for 20–30 min.

### Human PBMC isolation and CD14^+^ monocyte purification

Peripheral blood mononuclear cells (PBMCs) were isolated from buffy coats obtained from healthy donors by density gradient centrifugation using Ficoll-Paque PLUS. PBMCs were washed in PBS, filtered through a 40 µm strainer, and counted. For VLPs and EVs experiments, CD14^+^ monocytes were isolated using CD14 MicroBeads (Miltenyi Biotec) according to the manufacturer’s instructions, counted, and plated 75,000 cells/well in 96-well flat-bottom plates. For nucleofection experiments, CD14^+^ monocytes were isolated by negative selection using Pan Monocyte Isolation Kit, human (Miltenyi Biotec). For macrophage differentiation, monocytes were cultured in complete RPMI supplemented with recombinant human M-CSF (100 ng mL⁻¹) for 7 days (Peptrotech). To polarize MDMs, IFN-γ (20 ng/uL) and LPS (10 ng/uL) for M1, or IL-4 (20 ng/uL) for M2 were added at day 7. For dendritic cell differentiation, monocytes were cultured in complete RPMI supplemented with recombinant human GM-CSF (50 ng mL^-1^) (Miltenyi) and IL-4 (1000 IU mL^-1^) (ImmunoTools).

### Plasmid design and cloning

pCMV-MMLVgag-3xNES-Cas9 was purchased from Addgene n.181752 (Banskota et al., 2022). MLV packaging plasmids (Engels et al., 2022) and the VSV-G expression plasmid (Albanese et al., 2021) were available from our previous work. gRNAs targeting B2M were cloned into U6-driven backbone plasmid carrying the sgRNA scaffold. Briefly, gRNAs targeting B2M were synthesized as oligonucleotides and annealed in annealing buffer (10mM Tris pH8, 20mM NaCl) at 100 °C for 5 minutes followed by a slow cooling down to room temperature. The annealed oligonucleotides were cloned into a BsmBI digested U6-driven backbone plasmid using T4 DNA ligase (New England Biolabs) and transformed into TOP10 chemically competent E. coli. Correct clones were validated by Sanger sequencing (Eurofins Genomics), expanded, and plasmids were prepared using miniprep and maxi-prep kits (QIAGEN). List of primers and gRNAs is included as Supplementary Table.

### EVs and VLPs production

HEK293T cells were seeded 24 h before transfection to reach ∼60–70% confluence at transfection. Transfections were performed using linear polyethyleneimine (L-PEI; Polysciences) at a DNA:PEI ratio of 1:4 (µg:µl). DNA and PEI were separately diluted in serum-free DMEM, combined 1:1 (v/v), incubated for 20 min at room temperature to allow polyplex formation, and added directly to cells. After ≥4 h, medium was replaced with DMEM containing 2% FBS-EV depleted and cells were incubated for an additional 72 h before collection of conditioned medium. For large scale production, HEK293T were seeded in T175 flask format and transfected with 35 ug of total plasmid DNA. For VLPs production, plasmids encoding Cas9 fused to MLV Gag (20% of total plasmid DNA), MLV packaging functions (Gag-Pol) (20 % of total DNA plasmid), U6-driven gRNA expression (50% of total plasmid DNA), and VSV-G (at variable proportions) were co-transfected in HEK293T cells. For EV production, the packaging component was omitted (replaced with carrier DNA) while maintaining gRNA and VSV-G components. Conditioned medium was collected 72 h post-transfection, cleared by centrifugation (3,000g, 15 min), and filtered (0.45 µm, PES filters). Vesicles were purified either by PEG precipitation or ultracentrifugation depending on the application. For PEG precipitation, PEG solution was added to filtered supernatant at a 1:2 ratio (PEG:supernatant, v/v), mixed, incubated at 4 °C for 4 h, and pelleted (1,500g, 1 h, 4 °C). Pellets were resuspended in pre-filtered PBS, typically achieving ∼100-fold concentration, and stored at −80 °C. For ultracentrifugation, a sucrose cushion was layered into ultracentrifuge tubes prior to adding filtered supernatant. Samples were centrifuged at 100,000g for 2h at 4°C in a swinging bucket rotor (SW32, Beckman Coulter), supernatants were discarded, and pellets were resuspended in filtered PBS (typically ∼100-fold concentration). Aliquots were stored at −80 °C. **Nanoparticle tracking analysis (NTA)**

Particle size and concentration were measured using a NanoSight NS300 instrument. Vesicle preparations were diluted 1:100–1:1,000 in filtered PBS to a final concentration of 10⁷–10⁸ particles mL⁻¹. Three 30 s videos were acquired per sample (camera level 14–15) and analyzed with fixed gain settings; detection threshold was set between 3 and 5.

### Vesicle-mediated CRISPR/Cas9 editing

SUP-T1 cells (U-bottom 96-well plate) were treated with EVs or VLPs carrying Cas9 and one or two gRNAs targeting B2M. Cells were cultured for ≥3 days and split as needed to maintain optimal growth before knockout assessment by flow cytometry (typically day 5 post-treatment). Immediately after CD14 selection, monocytes (7.5 × 10⁴ per well, flat-bottom 96-well plate) were treated with EV/VLP dilutions. LPS (50 ng mL⁻¹) was used as a positive control for macrophage activation where indicated. For paired functional and viability measurements, parallel plates were prepared. The following day, supernatants were collected, cleared by centrifugation, stored at-80 °C for ELISA, and replaced with fresh RPMI supplemented with M-CSF. Cells were differentiated for 7 days prior to knockout validation. For moDC differentiation, monocytes were seeded as before in complete RPMI supplemented with GM-CSF and IL-4, treated with EV/VLPs dilutions and let differentiate for 7 days prior knockout validation.

### Nucleofection control (RNP delivery)

CRISPR RNP nucleofection was performed using a Lonza 4D-Nucleofector (program EH-100). Briefly, monocytes were washed, resuspended in P3 buffer, and mixed with pre-assembled RNP complexes generated by incubating sgRNA with NLS-Cas9 at room temperature for 15 min (Cas9:gRNA ratio 1:2.5; 40 pmol Cas9 per 100 pmol gRNA). Cells were recovered for 10 min in RPMI without supplements, then supplemented medium containing M-CSF was added and cells were plated as described above.

### Flow cytometry

For monocytes and SUP-T1, cells were stained with APC-conjugated anti-β2-microglobulin antibody (BioLegend, clone A17082A) diluted in FACS buffer and incubated for 20 min at 4 °C in the dark, then washed and analyzed on a BD FACSCanto II. For macrophages, cells were detached with accutase, incubated with Fc receptor blocking buffer (PBS with EDTA and 2% human serum), and stained with anti-β2-microglobulin APC antibody (clone A17082A) and a fixable viability dye (Near-IR) prior to acquisition. Where required, cells were fixed with 2% PFA. Data were analyzed in FlowJo (v10.10.0). For MDM, M1, M2, and moDC phenotype characterization, cells were detached and stained with the following antibodies: CD163 FITC (clone GHI/61), HLA-DR BUV805 (clone G46-6), HLA-ABC BUV 563 (clone W6-32), CD14 BB700 (clone M5E2), CD86 APC (clone IT2.2), CD40 BV605 (clone 5C3), CD206 BUV 395 (clone 19.2), CD209 Pe-Cy7 (clone 9E9A8), CD64 FITC (clone 10.1), CD11b BV421 (clone ICRF44), CD11c PE (clone B-ly6). For TLR4 and CCR5 detection, cells were detached as previously described and staining was performed at RT for 25 minutes using the following antibodies: TLR4-APC (clone HTA125) and CCR5 FITC (clone HEK/1/85a).

### Cell viability assay

Cell viability was assessed using CellTiter-Glo (Promega) according to the manufacturer’s protocol. Plates were incubated with reagent for 10 min in the dark with gentle shaking and luminescence was measured on a Synergy H1 plate reader (BioTek). Macrophage morphology was documented by brightfield microscopy (Zeiss Axio).

### Amplicon sequencing (MiSeq) and indel quantification

To quantify editing outcomes at the B2M locus, cells were collected 7 days post-treatment, washed and lysed in buffer containing CaCl₂, MgCl₂, EDTA, Triton X-100 and Tris (pH 7.5) supplemented with Proteinase K. Lysates were incubated at 65 °C for 10 min followed by 95 °C for 15 min and stored at −20 °C. The target locus was amplified by PCR using locus-specific primers carrying Illumina-compatible barcodes (PCR-I). PCR-I products were used as input for a second PCR (PCR-II) to add sample indices/adapters. Libraries were sequenced on an Illumina MiSeq platform and analyzed using the OutKnocker web tool. Indel frequency was calculated as:

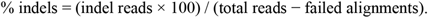

### ELISA

After 7 days from treatment, MDM NTC and MDM-TLR4 KO or untreated MDM were washed with PBS and stimulated with 50 ng/uL of LPS for 6 hours. Supernatant was recovered, clarified by ultracentrifugation at 2000 g for 10 minutes to remove any debris and tested for TNF-α and IL-6 using ELISA Flex kits (Mabtech) according to the manufacturer’s instructions with two modifications: incubation buffer contained 1% BSA (instead of 0.1%) and standards/samples were loaded at 60 µl per well (instead of 100 µl).

IFN-γ was measured using BD OptEIA™ Human IFN-γ ELISA Set according to the manufacturer’s instructions.

### Western blot

HEK293T cells and vesicle preparations were lysed in RIPA buffer supplemented with protease inhibitors. Lysates were clarified by centrifugation, and protein concentration was determined by BCA assay. Samples were denatured in LDS sample buffer, resolved on stain-free TGX gels (Bio-Rad), and transferred to nitrocellulose membranes using wet transfer. Membranes were blocked in 5% milk in TBS-T and probed with an anti–S. pyogenes Cas9 mouse mAb (Cell Signaling; clone 7A9-3A3), anti-Synthenin-1 rabbit mAb (Cell Signaling; clone E2I9L), anti-CD81 mouse mAb (ThermoScientific; clone M38), followed by HRP-conjugated anti-mouse secondary antibody. Signal was developed using ECL and imaged on an iBright 1500.

### Polyclonal alloreactive CD8 T cells

CD8 T cells were isolated from total PBMCs from one HLA-A *02:01 negative healthy donor using CD8 human MicroBeads (Miltenyi Biotec) according to the manufacturer’s instructions. CD8 T cells were seeded at a density of 1 million mL^-1^ in complete RPMI and repeatedly challenged with irradiated PBMCs from HLA-A *02:01 positive healthy donors to allow the selection of alloreactive CD8 T cells HLA-A *02:01 specific, supplementing them with IL-2 (50 IU/ml) only from the second re-stimulation. Polyclonal CD8 T cells were maintained in culture in complete RPMI supplemented with IL-2 (500 IU mL^1^). Cells were stimulated every 20 days with PHA (1 ug/mL) (Remel) and irradiated PBMCs pooled from four different donors.

### Killing and activation assays

Monocytes were isolated from an HLA-A*02:01 positive donors, seeded at 50,000 cells per well in 96 flat bottom plate in complete RPMI supplemented with M-CSF and treated with 15 x10^8^ VLPs and 60 x10^8^ EVs carrying NTC control gRNA or gRNAs 1 and 2 targeting B2M and let differentiate for 7 days. For proliferation and activation studies, after 7 days from MDM treatment, allogenic polyclonal CD8 T cells at resting state were washed with PBS and stained with Cell Trace Violet (CTV) (Invitrogen) according to manufacturer’s instruction. Stained CD8 T cells were counted and seeded in complete RPMI supplemented of m-CFS on top of treated MDM at a ratio of T:MDM of 0.5:1 and 1:1.

The following day, supernatants were collected, cleared by centrifugation, stored at −80 °C for ELISA for IFN-γ, and replaced with fresh RPMI supplemented with M-CSF. After 7 days, an image of each well was acquired using IncuCyte (Sartorius). Subsequently, T cells were resuspended, washed and stained with anti CD25-PE, anti CD8-APC and a fixable viability dye (Near-IR) and acquired at FACS Canto II at fixed time.

### Digital PCR for gRNA quantification

RNA was extracted from 50 uL of purified VLPs and EVs using QIAGEN miRNeasy kit following manufacturer instructions. Extracted RNA was quantified by Qubit™ RNA (Broad Range) (ThermoScientific), diluted 1×10^6^ and used as template for one-step digital PCR (QIAcuity One, 2plex Device) using QIAcuity OneStep Advanced EG Kit and QIAcuity Nanoplate 8.5k 24-well. Primers annealing to the sgRNA sequence were diluted to final concentration of 0.75 μM.

### VLPs vpx and HIV-1 production

VLPs, carrying Vpx from SIVmac251, were produced by co-transfection of 293T cells with pSIV3+ gag pol expression plasmids and the plasmid encoding VSV-G as previously described (Schneider et al., 2017). For HIV infection, the virus was produces as previously described (Albanese et al., 2024). HIV-1 GFP proviral clone NLENG1-I-7079 was used, referred to as R5 HIV-1 GFP in the current study. To produce sucrose cushion-purified HIV-1 stocks. 293T cells were seeded at a density of 8 x10^6^ cells in a 15-cm dish. After 24 h, cells were co-transfected with a mixture of 37.5 mg HIV-1 plasmid and 112.5 mL of L-PEI in DMEM without any additives for 30 min. After this time, the DNA/PEI solution was added on top of the cells. After 72 h, the supernatant was harvested and virus was purified via 25% sucrose-cushion centrifugation.

## Data analysis and statistics

Flow cytometry data were analyzed using FlowJo, exported, and plotted in GraphPad Prism (v10.4.1). For SUP-T1 knockout, values were normalized as:

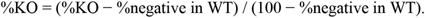

Statistical tests and replicate definitions are indicated in the figure legends.

## Data availability statements

The data supporting the findings of this study are provided in the Supplementary Information. Additional source data are available from the corresponding author upon reasonable request.

## Acknowledgements

We thank Chiara Angolini e Maria Cristina Crosti for the technical support. This work was supported by the Fondo Italiano Scienza (FIS2) from MUR (M.A.), and the MFAG AIRC (no. 28809 M.A.).

## AUTHOR CONTRIBUTIONS

Conceptualization: SF, MA

Methodology: SF, MR, AR, ANF, MA

Investigation: SF, MR, AR, ANF, TJNS, EP, ICS. PC, AM, ES, NAS

Visualization: SF, MR, MA

Supervision: GP, MA

Discussion: SF, GP, VH, TT, MA

Writing of original draft: SF, MR, NAS, VH, GP, MA

Review and editing: All authors.

