## Supplementary Info for "A vesicle-based platform for high-efficiency, high-viability CRISPR/Cas9 knockout in primary human myeloid cells"

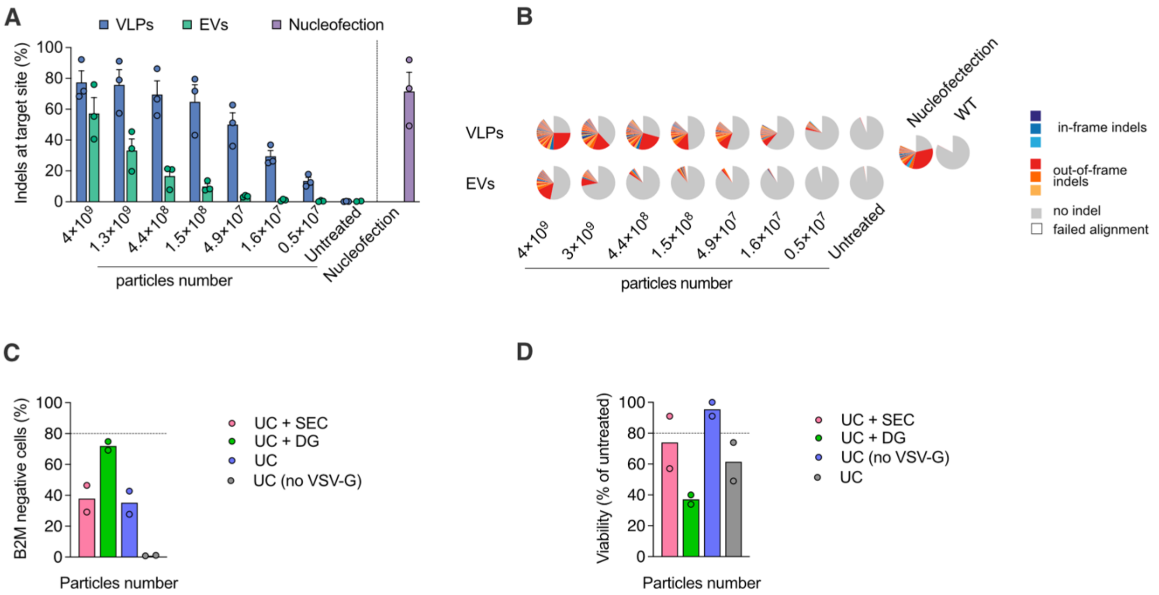


**Fig. S1 B2M editing efficiency and impact of purification on editing performance.**

Percentage of insertion and deletions (Indels) at the B2M locus in MDM treated with different concentrations of VLPs and EVs or nucleofected with RNP constituted of one gRNA. Cells were lysed after 7 days from treatment to perform PCR followed by Illumina Miseq sequencing. Analysis of reads was performed with Outknocker.2 web tool (mean ± SEM, n=3 donors). (**B**) Representative editing outcome plot from one donor generated using the OutKnocker web tool. Out-of-frame indels are shown in shades of red, and unedited reads (no indels) are shown in gray. (**C, D**) Comparison of VLP purification strategies: ultracentrifugation (UC), size exclusion chromatography (SEC), and density gradient (DG) and their impact on B2M editing (**C**) and viability (**D**) in primary $CD14^+$ monocytes treated with 5.56 $x10^8$ particles. B2M knockout efficiency was assessed by flow cytometry after differentiation. (mean ± SEM, n=2 donors)

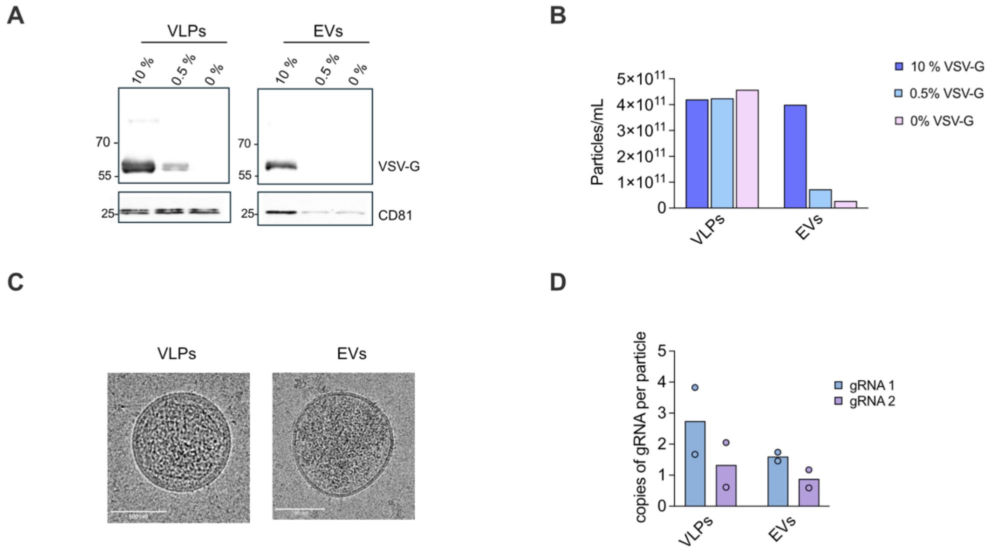


**Fig. S2. Characterization of EV and VLP production across VSV-G doses** (**A**) VSV-G detection in EVs and VLPs produced with high doses of VSV-G (10%), Low doses of VSV-G (0.5%) and no VSV-G by western blot. One representative blot is shown (n=2). (**B**) Particle concentration after ultracentrifugation-based purification of VLPs and EVs produced with high doses of VSV-G (10%), Low doses of VSV-G (0.5%) and no VSV-G measured at Nanosight 3000. (**C**) Cryo-electromycroscopy image of VLPs and EVs produced with MLV Gag-Cas9 and low doses of VSV-G. (**D**) Quantification of gRNAs 1 and 2 in EVs and VLPs measured by digital PCR. Copies of gRNA are normalized by particle number. (mean ± SEM, n=2 productions).

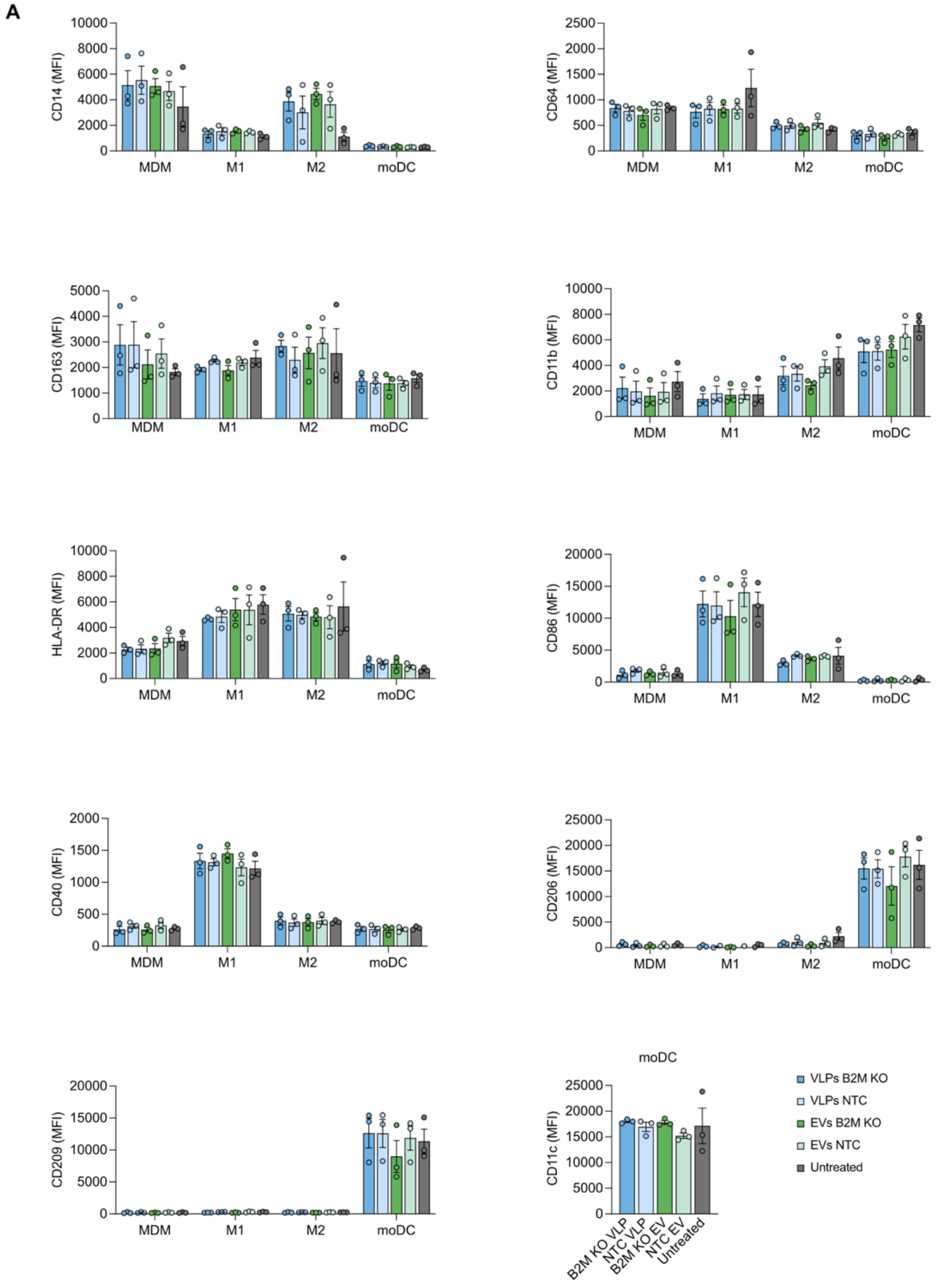


**Fig. S3. Differentiation makers evaluation in MDM, M1, M2, and moDC populations**

(**A**) Median fluorescent intensity (MFI) of surface markers that identifies MDM, M1, M2, and moDC populations. Monocytes were treated with VLPs and EVs targeting B2M or incorporating non-targeting control (NTC) gRNA followed by differentiation with different cytokine combinations (see material and methods).

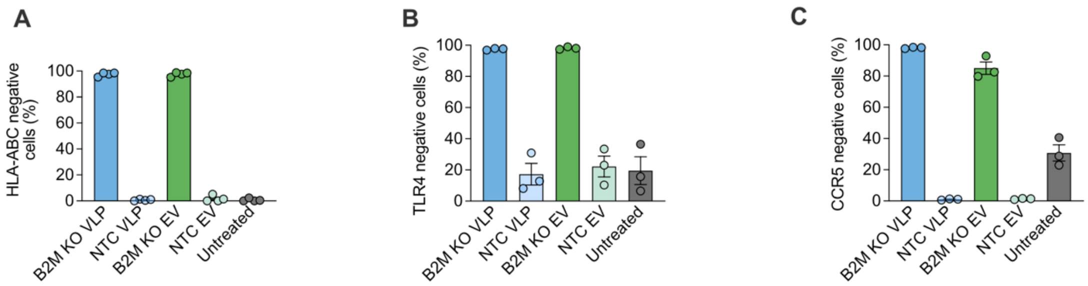


**Fig. S4: Evaluation of HLA-ABC, TLR4 and CCR5 knockout**

(**A**) Knockout efficiency of HLA-ABC in MDM treated with VLPs and EVs loaded with gRNA1 and gRNA2 targeting B2M locus. HLA-ABC expression was measured at FACS after 7 days from treatment. (mean ± SEM, n=3 donors). (**B**) Knockout efficiency of TLR4 in MDM treated with VLPs and EVs loaded with 2 gRNAs targeting TLR4 locus. TLR4 expression was measured at FACS after 7 days from treatment. (mean ± SEM, n=3 donors). (**C**) Knockout efficiency of CCR5 in MDM treated with VLPs and EVs loaded with 2 gRNAs targeting CCR5 locus. CCR5 expression was measured at FACS after 7 days from treatment. (mean ± SEM, n=3 donors).
